# Egg hatching success is significantly influenced by the time of thermal stress in multiple hard tick species

**DOI:** 10.1101/2022.12.13.518051

**Authors:** Oluwaseun M. Ajayi, Kennan J. Oyen, Benjamin Davies, Geoffrey Finch, Benjamin D. Piller, Alison A. Harmeyer, Katherine Wendeln, Carlie Perretta, Andrew J. Rosendale, Joshua B. Benoit

**Affiliations:** Department of Biological Sciences, University of Cincinnati, Cincinnati, OH, 45221, USA; Animal Diseases Research Unit, USDA-ARS, 3003 ADBF, Pullman, Washington, United States of America; Biology Department, Mount St. Joseph University, Cincinnati, OH, 45233, USA

**Keywords:** Thermal exposure, egg viability, ixodid ticks, embryo development, climate change

## Abstract

Ticks are blood-feeding arthropods responsible for the transmission of disease-causing pathogens to a wide range of vertebrate hosts, including livestock and humans. Tick-borne diseases have been implicated in significant economic losses to livestock production, and this threat will increase as these obligate parasites widen their geographical ranges. Just like in other ectotherms, thermal stress due to changing global temperatures has been shown to influence tick survival and distribution. However, studies on the influence of extreme temperatures in ticks have focused on advanced, mobile stages, ignoring stages that are immobile and cannot move to more favorable microhabitats. In this study, low- and high-temperature regimens were assessed in relation to egg viability for hard tick species - *Amblyomma maculatum* (Gulf Coast tick), *Ixodes scapularis* (black-legged tick), *Dermacentor variabilis* (American dog tick), and *Rhipicephalus sanguineus* (Brown dog tick). Tick eggs exposed early in development were significantly more susceptible to thermal stress when compared with those exposed later in development. In our tested models, treatment was more important for egg hatching than species differences. Lastly, there was evidence of extreme thermal exposure significantly altering the hatching times of tick eggs for specific treatments. These results provide insights into the critical period for tick egg viability and potential tick control strategies as the globe continues to experience climate change.

## Introduction

Thermal stress induced by changing global temperatures has been shown to have significant effects on many aspects of animal life including behavior (Kearney et al. 2009), physiological processes (Gunderson et al. 2016), and geographic distribution (Parmesan 2006). Due to global warming, terrestrial ectotherms like amphibians, non-avian reptiles, and invertebrates whose regulation of body temperature depends on external sources (Deutsch et al. 2008), are especially vulnerable to thermal stress. During the seasonal transitions of spring and fall, there are periods when temperatures fall far below freezing. Terrestrial arthropods tolerate this extreme cold through distinct physiological changes that range from increasing specific cryoprotective molecules that improve cold tolerance to changes in cell membranes that prevent membrane damage as the temperature lowers (Colinet et al. 2015, Overgaard and MacMillan 2017). Many arthropods enter a period of suspended development to survive through the winter (Denlinger 2022). Once winter ends, many arthropods lay eggs during early spring, which can experience freezing that could impact egg viability. As eggs cannot move into more favorable microhabitats, they are exceptionally vulnerable to local thermal changes (Pratt 1959, Zhang and Xin 1989, Anderson and Magnarelli 2008).

Thermal stress affects arthropods in many ways through its negative consequences on different biological parameters. The effects of heat and cold stresses include high mortality, reduced adult longevity, locomotor defects, decreased fecundity, impaired mating capacity, and biased sex ratios (Marshall and Sinclair 2010, Colinet et al. 2015, Jerbi-Elayed et al. 2015, Garcia and Teets 2019, Edmands 2021). Heat stress has been shown in male *Drosophila* to cause a reduction in sexual attractiveness (Fasolo and Krebs 2004), and impaired courtship owing to anatomical injuries (Rohmer et al. 2004, Krebs and Thompson 2005). Exposure to thermal stress in immobile stages (particularly the egg) is significant, resulting in detrimental outcomes such as the production of low-quality individuals (Sibly and Atkinson 1994), increased egg-larval mortality (Rocha et al. 2017, Zhou et al. 2018), and reduced larval growth (Potter et al. 2011). Several mite species, including ticks, are vulnerable to thermal stress during their mobile stages (Apanaskevich et al. 2013, Lu et al. 2014, Yuan et al. 2015).

Ticks are obligate, bloodsucking, ectoparasitic arthropods that transmit disease-causing pathogens to a wide range of hosts (Dantas-Torres et al. 2012, Pfäffle et al. 2013). They are of significant public health and economic importance as vectors of disease-causing pathogens with over 50,000 cases reported in 2019 in the USA (Dantas-Torres et al. 2012, CDC 2022). Global economic losses as a result of tick-borne diseases that affect livestock are estimated in the range of $20 to $30 USD billion annually (Lew-Tabor and Rodriguez Valle 2016), which is predicted to grow (Raghavan et al. 2019, Silatsa et al. 2019). Tick population and distribution are consistently impacted by host abundance/range shifts, habitat fragmentation, and micro- and macro-climate changes (Madison-Antenucci et al. 2020). Like other ectotherms, drastic temperature changes experienced by ticks have a considerable impact on their life history (Ogden and Lindsay 2016, Ogden et al. 2021). In specific, low and high temperatures have been shown to impact the mortality rates, questing behavior, duration, and rates of development and reproduction of ticks (Vail and Smith 2002, Ogden et al. 2004, Eisen et al. 2016, Ogden and Lindsay 2016, Benoit et al. 2021, Fieler et al. 2021).

There has been an emphasis on the effects of thermal stress on the larval, nymphal, and adult stages of ticks, with the mechanisms underlying cold tolerance explored significantly in these stages (Yu et al. 2014, Wang et al. 2017, Holmes et al. 2018, Agwunobi et al. 2021, Benoit et al. 2021, Fieler et al. 2021, Rosendale et al. 2022). However, there exists a dearth of research on thermal stress and tolerance limits in tick eggs. As tick eggs are sessile, the adult females determine where eggs will remain until larval emergence. Thermal stress has been shown to impact insect egg viability (Bowler and Terblanche 2008, Rocha et al. 2017), but many of these studies have been associated with high heat stress. Fed female ticks and larvae are observed throughout the year (Davidson et al. 1994, Kollars et al. 2000, Wilhelmsson et al. 2013, Fieler et al. 2021), indicating that eggs are likely exposed to both low- and high-temperature stress. In this study, we examined how different extreme low- and high-temperature regimens impact the egg viability of four Ixodid tick species: *Amblyomma maculatum* (Gulf Coast tick), *Ixodes scapularis* (black-legged tick), *Dermacentor variabilis* (American dog tick), and *Rhipicephalus sanguineus* (Brown dog tick). We compared differences in responses to cold and heat stress among species and the age (early and late development) of the tick eggs. Our results indicate that eggs exposed to thermal stress at an early stage have reduced larval emergence when compared to later stages. We also found out that treatment had a significant effect on egg viability with slight variations among the species, and the timing of egg hatching is influenced by extreme thermal exposure.

## Materials and methods

### Tick species

Engorged females (*A. maculatum*, *I. scapularis*, *D. variabilis*, and *R. sanguineus*) were obtained from the Oklahoma State University (OSU) Tick Rearing Facility (Stillwater, OK, USA). Tick colonies at this facility have been reared and often supplemented with field-collected individuals, and they are maintained at 22 ± 1°C, 93% relative humidity (RH), and 14:10 hr light: dark condition (L:D). Mated females were fed on sheep (*Ovis aries*). For our study, fed females were sent to our laboratory within 24 to 48 hrs of host drop-off. Upon arrival, fed females were held at 22 ± 1°C, 93% RH, and 14:10 hr L:D cycle (Winston and Bates 1960). As soon as the females laid eggs, some were collected immediately for early thermal exposure, while others required for late exposure were allowed to continue development. To prevent bias, eggs from multiple females (at least 8) were mixed before thermal stress protocols.

### Thermal stress experiments

#### General description of tick rearing and treatment

Early (freshly laid) and late eggs (∼2 weeks old identified with fecal spots; Figure 1A) of each tick species were subjected to different thermal stresses (described later). Based on a previous assessment of the specific stages of embryogenesis, the eggs were in stages 4-5 of development (embryonic cells have moved to the periphery of the egg and have begun to proliferate) for early thermal stress and stages 10-12 (presence of waste product in distinct sac-like structure and leg development can be observed) for late thermal stress (Dipeolu 1991, Santos et al. 2013). For each treatment, groups of 10 ticks (N = 10 groups) were placed in 1.5 cm^3^ Eppendorf tubes before being arranged into foam-plugged 50 mL Centrifuge tubes. Foam-plugged tubes were suspended in an ethylene glycol: water (60:40) solution whose temperature was regulated (±0.1°C) with a programmable bath (Arctic A25; Thermo Scientific, Pittsburgh, PA, USA), and relative humidity was maintained with a small vial of a supersaturated solution of potassium nitrate (93% RH). The rate of thermal change was ∼1°C per minute when the sample tubes were inserted into the water bath. Unless otherwise stated, eggs exposed to thermal stress (in 1.5 ml Eppendorf tubes with a mesh-covered hole to allow airflow) were then put in a mason jar containing a 25mL Erlenmeyer flask filled with water (∼20mL) and kept at rearing conditions previously described. Larval emergence was evaluated for all treatment groups at 2-week intervals. Control tick eggs were held at previously described rearing conditions but experienced no thermal exposure. The description of the different thermal stress regimes is as follows:

**Figure 1:**
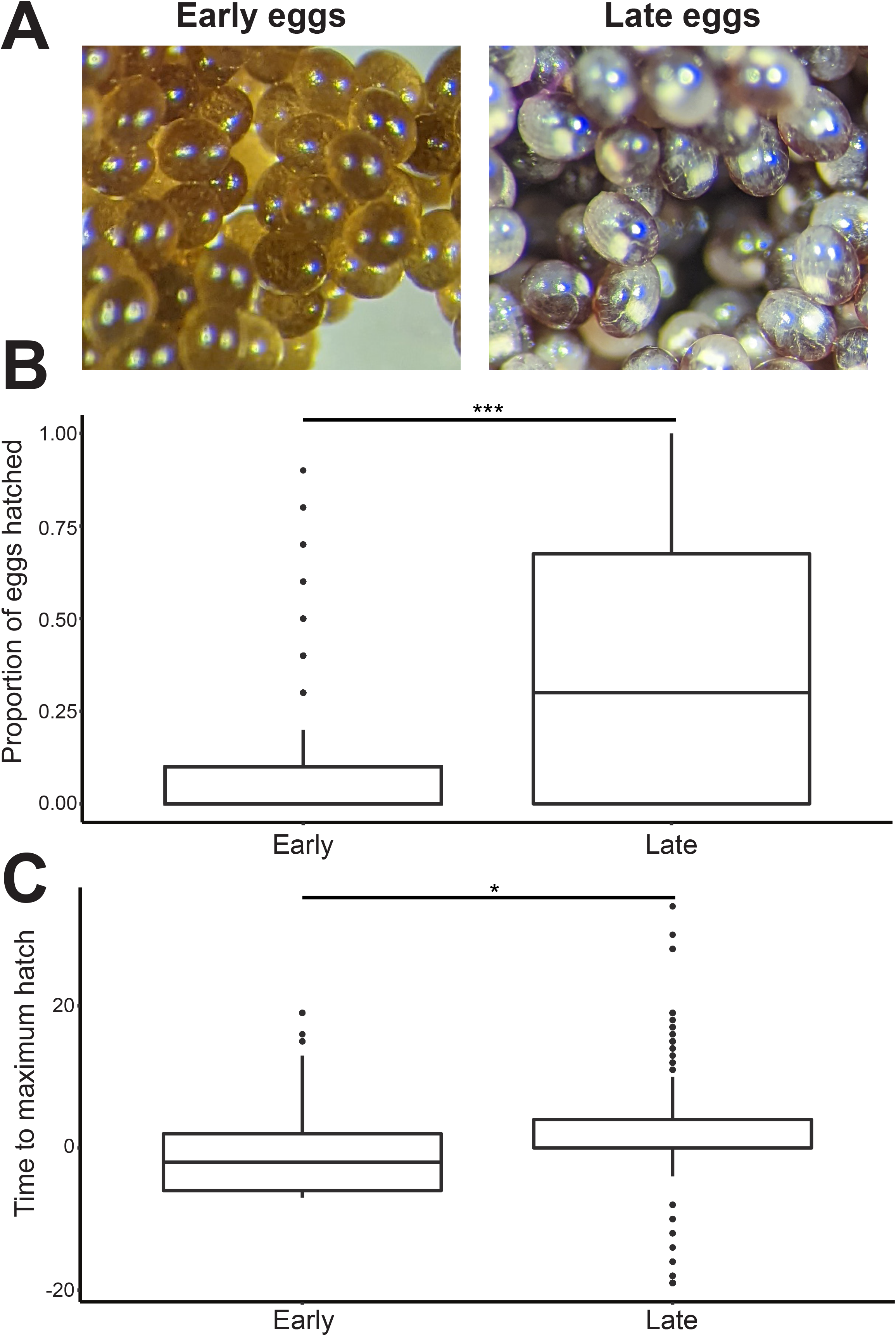
(A) Left: early tick eggs which are characterized by the movement of the embryonic cells to the periphery of the eggs and have begun to proliferate. Right: late tick eggs shown by the presence of waste product in a distinct sac-like structure and leg development can also be observed (Dipeolu 1991, Santos et al. 2013). Comparison of the (B) proportion of eggs hatched between early- and late-exposed eggs and C) residuals of time to maximum hatching between the early control and treatments and the late control and treatments pooled across all species. Symbols indicate statistical differences between groups (\**P*<0.05, \*\*\**P*<0.001). Summary statistics are found in Supplemental Materials S1 and S2. Ten groups of ten tick eggs were used for each treatment.

#### Cold shock

Tick eggs were placed in a bath for 2 hours at temperatures: −12.5, −15, −17.5, and −20°C (*I. scapularis*); −15, −17.5, −20, and −22.5°C (*A. maculatum*); −15, −17.5, and −20°C (*R. sanguineus*); and −15, −17.5, −20, −22.5, −25, and −27.5°C (*D. variabilis*). The variation in temperatures between species was based on a small-scale pilot study and on observations of larval cold tolerance (Fieler et al. 2021).

#### Cold acclimation for 2 hours and 7 days

To achieve 2 hours of acclimation, tick eggs were kept in an incubator at 0°C for 2 hours before exposure to low temperatures in the cold bath. A similar approach was used for 7 days of acclimation except that these eggs spent 7 days in the incubator. Temperatures tested were: −15, −17.5, and −20°C (*I. scapularis*); −17.5, −20, and −22.5°C (*A. maculatum*); −15, −17.5, and −20°C (*R. sanguineus*); and −17.5, −20, −22.5, −25, and −27.5°C (*D. variabilis*).

#### Inoculative freeze

Tick eggs were first put on ice in Eppendorf tubes before being placed in 50 mL tubes for the purpose of treatment in a cold bath at the following temperatures: −5, −10, −15, and −20°C (all the tick species). After 2 hours, the eggs were removed from the ice and placed in dry 1.5 ml Eppendorf tubes.

#### Heat shock

Tick eggs were placed in a warm bath at 42, 43, 44, and 45°C (all tick species) for 2 hours. The rate of thermal change was ∼1°C per minute.

#### Long-term (constant/cycle)

Tick eggs (in 1.5 ml Eppendorf tubes placed in a mason jar containing a 25mL Erlenmeyer flask filled with ∼20mL water) were placed at −5°C (constant) or cycled between −5°C and 3°C, to reflect the temperature range obtainable within a 24 h period in the tick microhabitat during mid-winter (Rosendale et al. 2016). The diurnal cycles in this period experienced 3°C for 7h and −5°C for 7h with a 1.6°C per hour cooling/warming between these temperatures. Tick eggs in the respective conditions were removed after 2, 4, 6, and 8 weeks.

### Data and statistical analyses

The total number of emerged larvae, the proportion of emerged larvae (emerged larvae/eggs treated), and the time to maximum hatching were collected for data analysis. Mean averages for all treatments are provided in Table S1 and S2. The proportion of eggs that hatched was calculated by dividing the number of hatched eggs by the total number of eggs for each treatment. The number of weeks to emergence was calculated by taking the number of weeks from the end of treatment until the last egg hatched and then averaging it across each vial and treatment. To account for the effect of early and late-stage eggs on the timing of egg hatching, we analyzed the differences between the early and late-stage treatments and subsequent controls, therefore early eggs were only compared with early controls, while late eggs were only compared with late controls. For the number of weeks till emergence, eggs that did not hatch were excluded from the analysis. We tested the effect of species, treatment, and developmental stage, on the proportion of eggs that hatched and the timing of hatching using a linear regression model, lm function in the lme4 R package. For the initial analysis of broad patterns within the data, we started with models with each response variable (time to maximum hatch and proportion hatched) as a function of each predictor including egg development stage, species, and treatment. For all linear models, a residual analysis plot (residual values vs. expected values) was visually examined to verify model assumptions of normality and homoscedasticity were met. Response variables were transformed (arcsin for proportion) as necessary to ensure that model assumptions are met. All additional analyses were performed using subsets of these models, including comparing within species, among treatments, and within development stages to allow for more targeted comparisons between treatments within species. Post-hoc comparisons were made using emmeans (default - Tukey method) where p < 0.05 were considered statistically significant. Output from models and post hoc results for these linear models are shown in Supplemental Materials S1 and Supplemental Materials S2. All analyses were done in R version 3.6.3 (statistical packages - linear model - lme4, with data, with posthoc - emmeans) and data visualization was completed using tidyr, ggplot, and reshape2.

## Results

### Exposure of eggs during early or late development alters the timing and proportion of egg hatching

The egg developmental stage during thermal stress significantly impacted the proportion of eggs that hatched and also the timing of hatching for pooled data of all treatments and species (Fig. 1B and 1C). Across all treatments and species, the proportion of eggs that hatched was significantly lower (F_1,2277_=150.6, *P*<0.00001) in the early-treated eggs compared with the late-treated eggs (Fig. 1B). For *D*. *variabilis*, about 25% more eggs were hatched in the late group compared with the early treated group (F_1,658_=159.9, *P*<0.00001). About 17% and 18% of eggs hatched more in the late groups of *R. sanguineus* (F_1,538_=29.11, *P*<0.00001) and *A. maculatum* (F_1,538_=50.42, *P*<0.00001), respectively, when compared with eggs treated early. But there was no significant difference between early- and late-exposed eggs of *I. scapularis* (F_1,537_=1.646, *P*=0.2).

In general, the egg developmental stage during stress exposure had a slight impact on the timing of hatching (Fig. 1C; F_1,1224_=4.545, *P*=0.034), where eggs exposed during the early stage tended to hatch slightly sooner than expected relative to the control group (with residuals falling below the zero line but this observation is not statistically significant); and those exposed during the late stage hatched significantly later than expected in respect to the control (residuals significantly above the zero line for some treatments). This effect was specifically driven by the cold-acclimated and cold-shocked eggs hatching sooner than expected in the early group, and cold acclimated and inoculative freezing treatments hatching later than expected in the late group. Early-stage *A. maculatum* control eggs hatched after 8.0 weeks while late-stage eggs hatched in 7.8 weeks. For *D. variabilis*, early-stage eggs hatched after 6.4 weeks and late-stage eggs hatched after 8.1 weeks. Early *I. scapularis* eggs hatched after approximately 7.0 weeks while late-stage eggs hatched after 7.7 weeks. For *R. sanguineus*, the late-stage eggs hatched after 9.7 weeks, while the early-stage eggs hatched after 9.1 weeks.

### Stressful temperature exposure has a differing effect on the proportion of eggs hatching for each species

In general, treatment had a significant effect on the proportion of eggs that hatched in all species. For *D. variabilis*, treatment had an overall significant influence on the proportion of eggs that hatched (Fig. 2A, F_32,627_= 17.37, *P*<0.00001). All temperature treatments had a deleterious effect on egg hatching for *D. variabilis*, significantly reducing the proportion of eggs that hatched compared with controls (Fig 2A). This pattern also held in *R. sanguineus* where many of the temperature treatments had little or no egg hatching unlike the control (Fig 2B, F_26,513_=37.68, *P*<0.00001). In *I. scapularis*, treatment also showed a significant influence on the proportion of eggs that hatched (Fig. 2C, F_26,512_=36.63, *P*<0.00001), with the strongest negative effects in the eggs exposed to heat shock, which was also the case for *R. sanguineus* and *D. variabilis*, with almost no eggs hatching in groups treated at such high temperatures. *A. maculatum* was an exception to this trend, with heat shock slightly increasing hatching proportions in the late-stage eggs (Fig. 2D, F_53,486_=25.71, all *P*<0.00001). Overall, eggs were more sensitive to heat exposure compared with cold exposures which either increased hatching proportions or had no effect.

**Figure 2:**
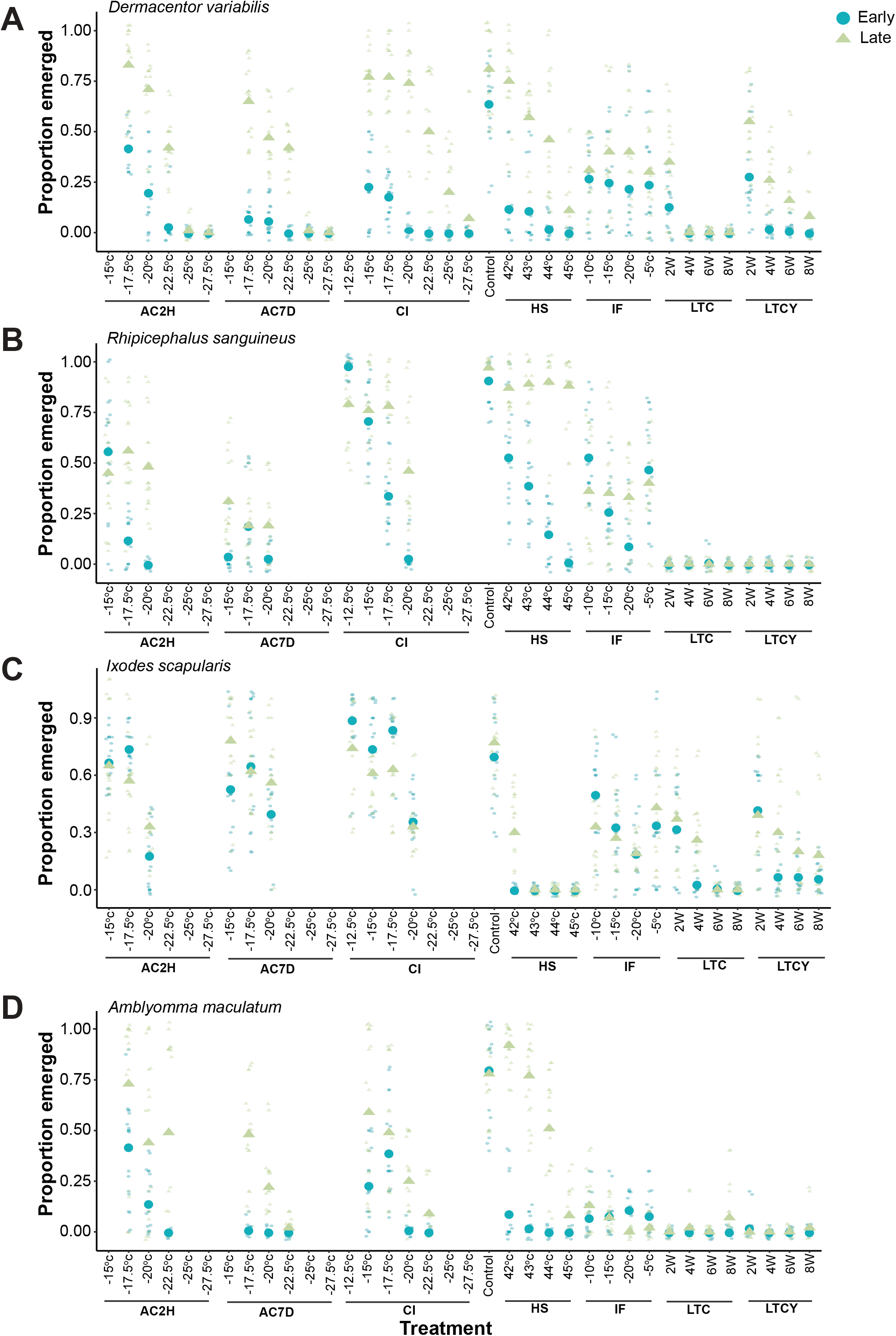
Comparison of proportion of larvae that emerged from both early and late eggs for all indicated treatments in (A) *Dermacentor variabilis*, (B) *Rhipicephalus sanguineus*, (C) *Ixodes scapularis*, and (D) *Amblyomma maculatum*. The treatment groups are as follows: AC2H, Cold acclimation for 2 hours; AC7D, Cold acclimation for 7 days; CI, Cold immediate; HS, Heat shock; IF, Inoculative freeze; LTC, Long term constant; LTCY, Long term cycle; and Control. Summary statistics are found in Supplemental Materials S1 and S2. Ten groups of ten tick eggs were used for each treatment.

### Differing thermal treatments alter the timing of egg hatching depending on the species

To ascertain the influence of stressful thermal exposures on the hatching duration of early and late tick eggs, we analyzed the differences in the number of weeks to larval emergence between the treatments and controls. Across all species, very few larvae emerged in cyclical cold treatments or constant cold treatments regardless of development stage and these were therefore excluded from further analysis. For *D. variabilis*, the only non-control group in the early-stage eggs that hatched were those that experienced cold acclimation for 2hrs before harsh cold exposure at −17.5°C and the time to emergence was not significantly different from the control (Fig. 3A, F_18,95_=1.089, *P*>0.05). Within the late-stage eggs, the only significant difference from the control was in the eggs treated at −20°C after 2 hours of acclimation, which hatched approximately 2 weeks earlier than the control group (Figure 3A). Although insignificant, several treatments in the late-stage eggs expedited larval emergence in comparison with the single treatment in the early-stage eggs which delayed emergence. Early-stage eggs exposed to heat shock at 42°C and 43°C significantly decreased the time to emergence by 2 weeks in *R. sanguineus*, as well as eggs that experienced an immediate cold exposure at −12.5°C (Fig. 3B, F_17,100_=3.25, all *P*<0.05). In the late-stage eggs, exposure to extreme cold at −15°C and −17.5°C significantly delayed larval emergence by 4 and 6 weeks respectively (Fig. 3B, F_18,160_=4.743, all *P*<0.02). In *I. scapularis*, treatment did not significantly affect the emergence timing in early-stage eggs (Fig. 3C, F_21,148_=1.282, all *P*>0.05), mainly driven by the very low number of early-stage eggs that successfully hatched. Of the eggs that did hatch, acclimation to cold temperatures tended to increase the time to larval emergence. In the late-stage eggs, cold exposure in the form of acclimation, inoculative freezing, and cold shock all significantly delayed larval emergence (Fig. 3C, F_21,181_=2.607, all *P*<0.05). All models of *A. maculatum* egg hatching duration were insignificant, due to the low numbers of larvae that successfully emerged. Of those that did emerge, both warm and cold exposure tended to increase the time to egg hatching. In general, cold treatments delayed egg hatching and warm treatments tended not to affect egg hatching duration.

**Figure 3:**
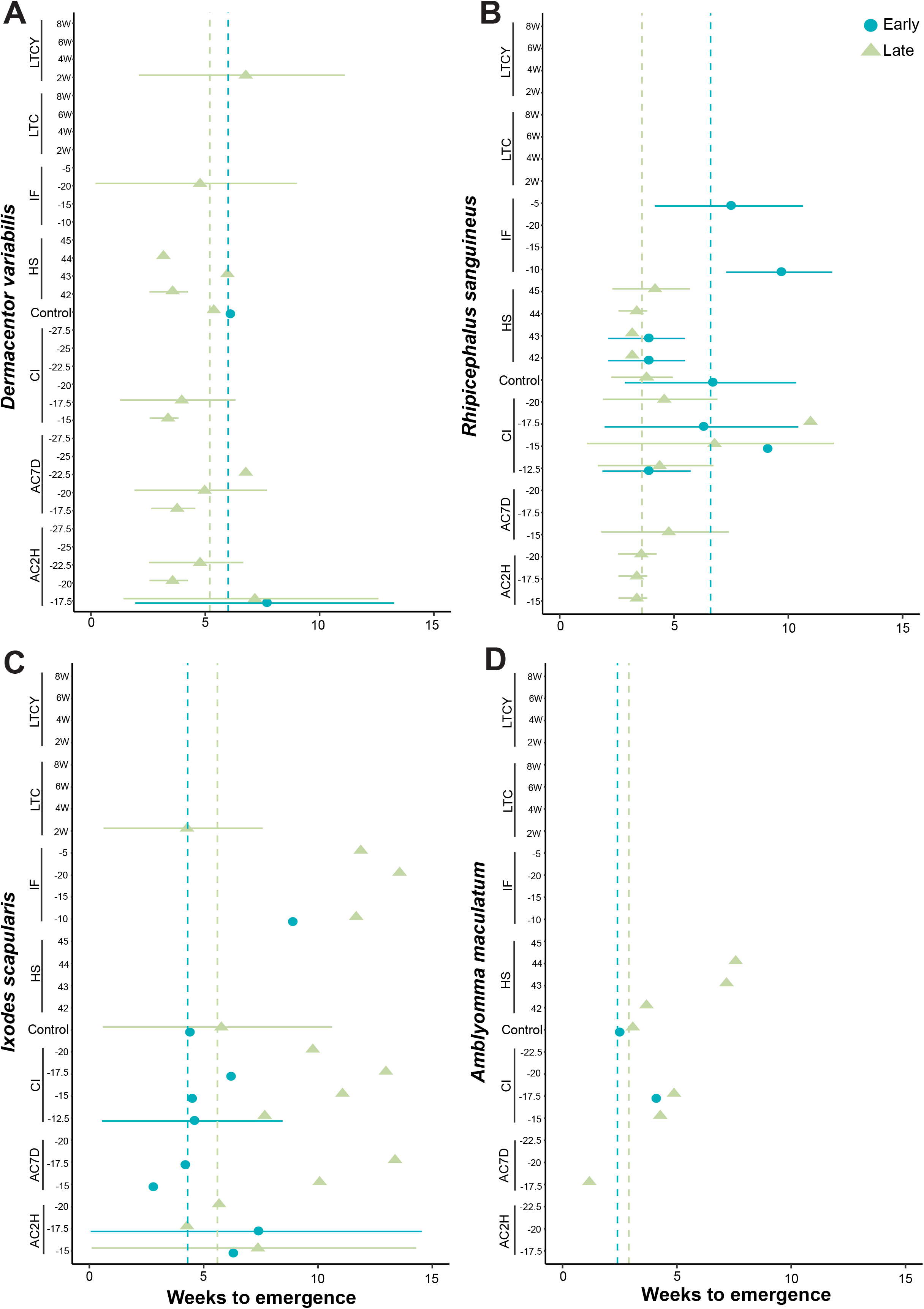
Comparison of weeks to larval emergence from both early and late eggs for all indicated treatments in (A) *Dermacentor variabilis*, (B) *Rhipicephalus sanguineus*, (C) *Ixodes scapularis*, and (D) *Amblyomma maculatum*. The treatment groups are as follows: AC2H, Cold acclimation for 2 hours; AC7D, Cold acclimation for 7 days; CI, Cold immediate; HS, Heat shock; IF, Inoculative freeze; LTC, Long term constant; LTCY, Long term cycle; and Control. Dashed lines represent values for the control groups. Summary statistics are found in Supplemental Materials S1 and S2. Ten groups of ten tick eggs were used for each treatment.

### Acclimation influences the proportion and timing of egg hatching in late-exposed eggs

Heat shock did not significantly impact the timing of egg hatching in late-exposed tick eggs that survived such exposure across any species (Fig. 4A, F_3,6_=0.3053, all *P*>0.05). Cold acclimation had a significant impact on both the timing of egg emergence and the proportion of emerged eggs in those that received thermal exposure in the late stage (Figure 4B and 4C), where unacclimated eggs took ∼2.6 weeks longer to hatch than acclimated eggs after cold shock, (F_1,27_=4.921, all *P*=0.0351). For *R. sanguineus*, a greater portion of acclimated eggs hatched compared with the unacclimated eggs in both the short (hours) and long (days) term acclimation treatments (Fig. 4B, F_9,12_=3.689, all *P*<0.005). For *D. variabilis*, only the group acclimated for 7 days before cold exposure at −20°C was significantly different from unacclimated eggs (Fig. 4B, F_9,12_=3.689, *P*=0.0315). Although heat shock had little impact on the timing of egg hatching, cold shock tended to increase the development time of eggs, but acclimation at 0°C for 2 hours before cold exposure reduced this effect.

**Figure 4:**
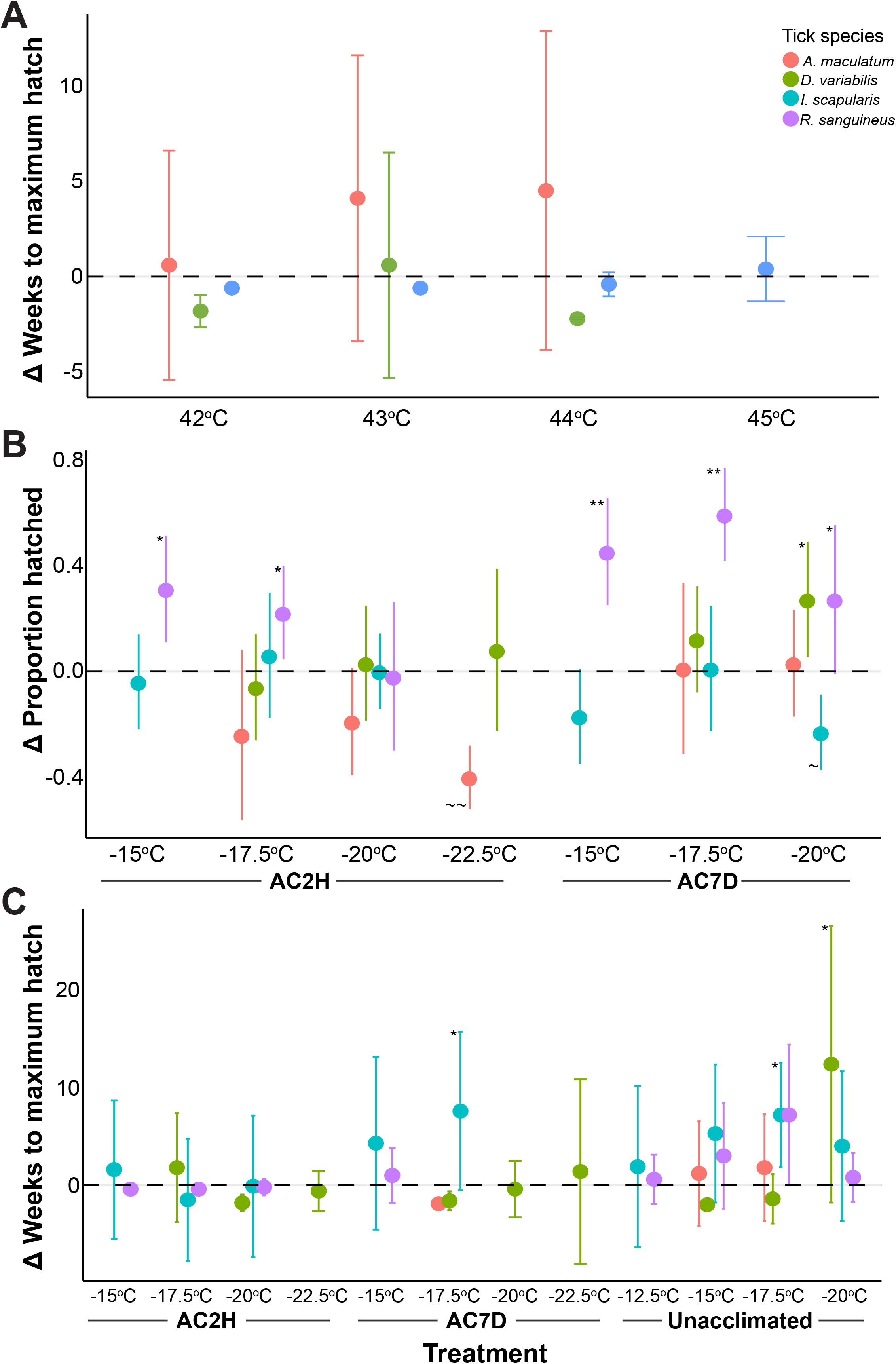
(A) Differences in the weeks to maximum larval emergence in the heat shock treatment among the multiple tick species in late-stage exposure. (B) Differences in the proportion of eggs that hatched in the cold acclimation treatment groups among the multiple tick species in late-stage exposure. The y-axis (Δ Proportion hatched) represents the proportion hatched in the cold acclimation minus the expected proportion hatched in cold shock. (C) Differences in the weeks to maximum larval emergence in the cold acclimation and cold shock treatment groups among the multiple tick species in late-stage exposure. The y-axis (Δ Weeks to maximum hatch) means the change in weeks that it took to reach maximum hatch in the cold-acclimated ticks compared with unacclimated ticks. Symbols indicate statistical differences from the zero line (*= positive direction, ∼ = negative direction; * and ∼ = *P*<0.05, ** and ∼∼ = *P*<0.01). AC2H and AC7D are cold acclimations for 2 hours and 7 days, respectively, while unacclimated represents the cold shock treatment. Summary statistics are found in Supplemental Materials S1 and S2. Ten groups of ten tick eggs were used for each treatment.

## Discussion

The proportion and timing of hatching were influenced by stressful temperature exposures of tick eggs in both early- and late-stage embryogenesis. Embryos are more sensitive to thermal stress than other life stages due to their relatively narrow thermal tolerance breadth and inability to behaviorally thermoregulate (Hallman and Denlinger 1998). Of importance, thermal tolerance measurements in eggs have been understudied when compared to other developmental stages (Bowler and Terblanche 2008, Tippelt et al. 2020, Borda et al. 2021, Kramer et al. 2021, Weaving et al. 2022). For tick eggs where larval emergence still occurred after thermal stress, we observed differences in developmental rates depending on when eggs were exposed. Specifically, exposure to both warm and cold temperatures in both early- and late-stage eggs delayed hatching and decreased the number of eggs that survived. Here, fewer than 40% of eggs treated during the early phase hatched successfully. This likely indicates that early embryonic stages are highly sensitive to thermal stress, which matches the results of other invertebrate systems (Hallman and Denlinger 1998, Pörtner et al. 2017, Harvey et al. 2020).

The developmental stage of the eggs strongly influenced responses to temperature, but this has been assessed in a few animals with a focus on heat stress (Graziosi et al. 1980, Harvey et al. 2020). We found that exposure during early embryogenesis was more deleterious than in the late embryogenesis stages in all species. This is consistent with thermal tolerance in vertebrates where major structural and organ systems rapidly develop in early embryonic stages and are more sensitive to environmental perturbations compared with later stages (Sadler 2018). Thermal stress in other terrestrial arthropods, such as in *Drosophila*, has indicated that embryogenesis is a period of low thermal resistance (Hallman and Denlinger 1998, Bowler and Terblanche 2008, Rocha et al. 2017, Harvey et al. 2020, Borda et al. 2021), but no studies have compared differences throughout embryonic development.

Most eggs did not survive heat shock exposure, but of those that did, heat shock did not significantly impact the timing of egg hatching. Significantly more eggs survived cold shock matching previous work on mobile stages that suggests that high-temperature exposures are more deleterious for ticks than low temperature exposures (Holmes et al. 2018, Ogden et al. 2021). Acclimated eggs tended to hatch sooner than unacclimated eggs following cold exposure (specifically in late-stage eggs), potentially suggesting an acclimation response in tick eggs at a later stage of development. Investigation in the mosquito, *Aedes albopictus* revealed that cold acclimation alters egg structure, a potential strategy for preventing ice formations (Kreß et al. 2016). It might be interesting to evaluate the egg ultrastructure of hard ticks in late embryogenesis to confirm if there is a similar mechanism of cold acclimation.

Interestingly, for *I. scapularis* and *D. variabilis* long-term cold exposures were more deleterious than exposures that cycled and only held at the coldest temperature for a few hours each day. Studies comparing fluctuating and constant temperature exposures have shown that fluctuating temperatures could enhance or weaken performance depending on the severity of the exposure (Colinet et al. 2015). Fluctuating exposures that repeatedly reach stressful high and low temperatures may have a cumulatively negative impact, while those that remain within permissible ranges might allow for repair from extreme exposures during the intermediate ranges (Kingsolver et al. 2015). Recent studies have shown repair mechanisms can reestablish system-wide homeostasis during fluctuating temperature regimes through the production of heat shock proteins, or low molecular weight solutes (Colinet et al. 2018). For *A. maculatum* and *R. sanguineus*, −5°C was likely outside of the tolerable limit as they experienced high mortality in both fluctuating or constant conditions. This hints at an enhanced capacity for eggs of *D. variabilis* and *I. scapularis* to withstand cooler conditions and potentially repair damage induced by stressful temperature exposure compared with *A. maculatum* and *R. sanguineus*.

Stressful temperature exposure altered the time to hatching for both early- and late-stage eggs, although the effects were weak for early-stage eggs, due to the limited number of eggs that survived. For eggs exposed to heat shock, there were no significant differences between the treatments and controls, but some tended to hatch earlier than controls. This matched what we expected because exposure to warmer temperatures typically increases growth and development in arthropods (Cossins and Bowler 1987, Rueda and Axtell 1996). Interestingly, there was no significant difference between cold and warm exposure in either the timing of development or survival among the early-stage eggs, suggesting that during this short time window, tick eggs are extremely vulnerable to any suboptimal temperature exposure. *D. variabilis* was an exception to this pattern with several late-stage eggs hatching earlier than predicted across several treatments of both cold and hot temperatures. Of all the species tested, *D. variabilis* had the greatest proportion of late-stage eggs that emerged following treatments. Studies on comparative tick larval thermal tolerance show that the American dog tick is the most cold-tolerant tick species (Fieler et al. 2021), which we now observe in the egg stage. Why cold temperature exposure increased the rate of development in some *D. variabilis* eggs is an open question. Still, it could be related to stressful temperature exposure increasing egg metabolic rates, leading to faster growth and development.

Generally, eggs of *D. variabilis* were more robust to thermal stress than those of the other three species. This matches our prediction, given the thermal variability hypothesis, which suggests that broadly distributed species should have comparably wide thermal tolerance ranges (Stevens 1989). Of the tested species, *D. variabilis* is the most broadly distributed (Minigan et al. 2018) and has eggs with correspondingly wide thermal tolerance ranges. In contrast, *A. maculatum*, which is generally restricted to southern portions of North America, has the lowest tolerance to stressful temperatures and *I. scapularis* the second lowest, which is also narrowly distributed compared with *D. variabilis* and *R. sanguineus*. Our findings are consistent with the climatic variability hypothesis, with the most broadly distributed ticks species being more robust to temperature variability, while narrow thermal tolerance was correlated to narrow geographic distributions. Although the climate variability hypothesis has been extensively tested in arthropods (Sunday et al. 2011), studies on eggs have been limited. Investigations using both tropical and European eggs of *Aedes aegypti* and *Ae. albopictus* revealed that differences in hatching success between species within the same geographical range were less significant than differences between geographically different strains of the same species (Thomas et al. 2012). Also, other factors could be responsible for species differences in thermal tolerance. For example, a comparison of cold tolerance of both *Ae. aegypti* and *Ae. albopictus* show that cold winter does not seem to be the factor restricting the distribution of *Ae. aegypti* to subtropical/tropical habitats, since their eggs can survive at −2°C for 6 days (Kramer et al. 2020). The inability of *Ae. aegypti* to induce dormancy (at the egg stage), just like what is seen in *Ae. albopictus* might explain their inability to persist in colder climates.

Although several studies have examined the thermal tolerance of ticks during active life stages (Rosendale et al. 2016, 2022, Wang et al. 2017, Holmes et al. 2018, Fieler et al. 2021), little is known about egg tolerance. Our study addresses this by providing a more holistic understanding of how early developmental stages influence tick populations and how ticks will respond to shifts in environmental conditions. Models that assess the potential for ticks to shift their ranges to new environments may overestimate their results given the relatively narrow thermal tolerances of tick eggs compared with active life stages. Additionally, the initial choice of where the mothers deposit their eggs will have considerable importance in tick egg viability in relation to thermal exposure. Importantly, thermal tolerance will likely need to be examined for ticks collected from multiple geographic regions as significant differences in thermal tolerance has been noted in other terrestrial invertebrates among population (Arthur et al. 2008, Sinclair et al. 2012, Clemson et al. 2016) Successful embryonic development is critical for survival in every environment and should be considered when assessing the impacts of climate change on populations and when devising tick control strategies. Importantly, as the thermal tolerance of eggs during embryogenesis has not been extensively investigated, similar effects are likely to occur for other arthropod species.

## Acknowledgments

This work was supported by the United States Department of Agriculture’s National Institute of Food and Agriculture Grant 2016-67012-24652 to A.J.R. Funding was also provided by the University of Cincinnati as a Faculty Development Research Grant and, in part for dual usage of equipment, through the National Science Foundation DEB-1654417 and National Institute of Allergy and Infectious Diseases of the National Institutes of Health under Award Number R01AI148551 to J.B.B. K.J.O was funded by the David H. Smith Conservation Fellowship.

## Supplemental resources

**Table S1:** Mean proportion of eggs hatched for each treatment group, developmental stage, and species.

**Table S2:** Mean timing of eggs hatched for each treatment group, developmental stage, and species.

**Supplemental materials 1:** Global model (generalized linear) results for the effect of treatment, developmental stage, and species on the proportion of eggs that hatched.

**Supplemental materials 2:** Global model (generalized linear) results for the effect of treatment, development stage, and species on the timing of egg hatching.

